# The domain-general multiple demand (MD) network does not support core aspects of language comprehension: a large-scale fMRI investigation

**DOI:** 10.1101/744094

**Authors:** Evgeniia Diachek, Idan Blank, Matthew Siegelman, Josef Affourtit, Evelina Fedorenko

**Author notes:** Corresponding Authors Evgeniia Diachek and Ev Fedorenko,; 43 Vassar Street, Room 46-3037, Cambridge, MA, 02139. Equal contributors (Evgeniia Diachek, Idan Blank, Matthew Siegelman).

## Abstract

Aside from the language-selective left-lateralized fronto-temporal network, language comprehension sometimes additionally recruits a domain-general bilateral fronto-parietal network implicated in executive functions: the multiple demand (MD) network. However, the nature of the MD network’s contributions to language comprehension remains debated. To illuminate the role of this network in language processing, we conducted a large-scale fMRI investigation using data from 30 diverse word and sentence comprehension experiments (481 unique participants, 678 scanning sessions). In line with prior findings, the MD network was active during many language tasks. Moreover, similar to the language-selective network, which is robustly lateralized to the left hemisphere, these responses were stronger in the left-hemisphere MD regions. However, in stark contrast with the language-selective network, the MD network responded more strongly (i) to lists of unconnected words than to sentences, and critically, (ii) in paradigms with an explicit task compared to passive comprehension paradigms. In fact, many passive comprehension tasks failed to elicit a response above the fixation baseline in the MD network, in contrast to strong responses in the language-selective network. In tandem, these results argue against a role for the MD network in core aspects of sentence comprehension like inhibiting irrelevant meanings or parses, keeping intermediate representations active in working memory, or predicting upcoming words or structures. These results align with recent evidence of relatively poor tracking of the linguistic signal by the MD regions during naturalistic comprehension, and instead suggest that the MD network’s engagement during language processing likely reflects effort associated with extraneous task demands.

**Significance Statement:** Domain-general executive processes, like working memory and cognitive control, have long been implicated in language comprehension, including in neuroimaging studies that have reported activation in domain-general multiple demand (MD) regions for linguistic manipulations. However, much prior evidence has come from paradigms where language interpretation is accompanied by extraneous tasks. Using a large fMRI dataset (30 experiments/481 participants/678 sessions), we demonstrate that MD regions are engaged during language comprehension in the presence of task demands, but not during passive reading/listening—conditions that strongly activate the fronto-temporal language network. These results present a fundamental challenge to proposals whereby linguistic computations, like inhibiting irrelevant meanings, keeping representations active in working memory, or predicting upcoming elements, draw on domain-general executive resources.

## Introduction

Converging evidence from neuroimaging and patient studies suggests that a left-lateralized fronto-temporal brain network is selective for language processing. These regions respond to linguistic input (visual or auditory) across diverse materials and tasks (e.g., Fedorenko et al., 2010, 2016; Vagharchakian et al., 2012; Scott et al., 2017; Deniz et al., 2019), but not to non-linguistic cognitive tasks, like arithmetic calculations, executive function tasks, music perception, action/gesture observation, and non-verbal social information (e.g., Fedorenko et al., 2011; Monti et al., 2012; Pritchett et al., 2018; Jouravlev et al., 2019; Paunov, 2019; see Fedorenko & Varley, 2016, for a review).

In addition to this “core” language-selective network, numerous fMRI language studies have reported activation in what-appear-to-be regions of a different network: a domain-general bilateral network of frontal, parietal, cingular, and opercular brain regions known as the multiple demand (MD) network (Duncan, 2010, 2013). This network supports diverse cognitive tasks (Duncan & Owen, 2000; Fedorenko et al., 2013; Hugdahl et al., 2015) and has been linked to constructs like working memory, cognitive control, and goal-directed behavior (Cole & Shneider, 2007; Duncan, 2010). The MD network is dissociated from the language network (see Fedorenko & Blank, 2020, for a review), as evidenced by brain imaging studies (Fedorenko et al., 2011; Blank et al., 2014; Mineroff et al., 2018), patient investigations (Woolgar et al., 2018), and gene expression patterns (Kong et al., 2020). Therefore, the two networks likely serve separable computational goals. However, many complex cognitive processes may rely on multiple distinct, and possibly interacting, cognitive mechanisms and their associated brain networks (Petersen & Sporns, 2015). Language comprehension may thus be supported by both the language-selective network and the domain-general MD network.

MD network engagement has been reported for diverse linguistic phenomena, including lexical/structural/referential ambiguity (e.g., Rodd et al., 2005; Novais-Santos et al., 2007; January et al., 2009; McMillan et al., 2013), high surprisal (e.g., Strijkers et al., 2019; cf. Shain, Blank, et al., 2020) including grammatical violations (e.g., Kuperberg et al., 2003; Nieuwland et al., 2012; Mollica et al., 2020), and syntactic complexity in unambiguous structures (e.g., Peelle et al., 2010). These results align with behavioral evidence for the role of working memory/cognitive control in language comprehension (e.g., King & Just, 1991; Gernsbacher, 1993; Waters and Caplan, 1996; Gibson, 1998; Gordon et al., 2002; Fedorenko et al., 2006, 2007; Lewis et al., 2006; Novick et al., 2009). Some have therefore proposed that domain-general executive resources— implemented in the MD network—support *core aspects of linguistic interpretation* related to lexical access, syntactic parsing, or semantic composition (Hasson et al., 2018), like inhibiting irrelevant meanings/parses (Novick et al., 2005), selecting the relevant representation from among alternatives (Thompson-Schill et al., 2002; Hirshorn & Thompson-Schill, 2006; Grindrod et al., 2008), supporting predictive coding for language processing (Strijkers et al., 2019), or keeping linguistic representations active in working memory (Moser et al., 2007).

Others, however, have questioned the importance of domain-general executive resources / the MD network to language processing (for reviews, see Fedorenko, 2014; Campbell & Tyler, 2018). For example, Wright et al. (2011) showed that some frontal regions— plausibly MD areas—are only engaged during a lexical decision task, but not passive listening to the same materials. And Blank & Fedorenko (2017) demonstrated that MD regions do not closely track the linguistic signal during comprehension of naturalistic stories, suggesting that they are unlikely to support computations that relate to the properties of the input (see also Wehbe et al., submitted).

To illuminate the role of the MD network in language processing, we conducted a large-scale investigation of diverse comprehension tasks. In particular, we used data from thirty fMRI experiments to examine the responses of MD and language regions—functionally defined in each participant using independent localizer paradigms (Fedorenko et al., 2010, 2013)—to different linguistic stimuli and tasks. To foreshadow the key results, we found above-baseline responses in the MD network during many linguistic tasks. However, passive comprehension tasks, which robustly engage the language-selective network, elicited a response at the level of the fixation baseline in the MD network. These results argue against the role of the MD network in core aspects of sentence comprehension, at least across the materials tested here and in neurotypical young adults.

## Materials and Methods

Because prior literature has not delivered a clear answer as to the role of the MD network (also sometimes referred to as the “executive/cognitive control network” or “task positive network”) in language comprehension, we here combined data from numerous diverse word and sentence comprehension experiments that have been conducted in our lab over the last decade. Given that each participant performed functional localizer tasks (e.g., Saxe et al., 2006) for the MD (and language) network, we could straightforwardly combine data from across experiments by pooling responses from functionally defined MD (or language) areas and have greater confidence that these constitute the ‘same’ regions (i.e., functional units) across individuals compared to relying on anatomy alone (e.g., Brett et al., 2002; Saxe et al., 2006; Fedorenko & Kanwisher, 2009; Fedorenko et al., 2010; Nieto-Castañon & Fedorenko, 2012; Fedorenko & Blank, 2020). The fact that the linguistic experiments varied in the presence of an explicit task (13 passive reading/listening experiments, 17 experiments with a task)—with the task further varying across experiments—allowed us to test the critical question of whether the MD network’s engagement is restricted to cases where an explicit task is present. (Note that a traditional whole-brain group-analytic approach (e.g., Holmes & Friston, 1998) would be unlikely to yield a clear answer in this study due to the combination of i) high inter-individual variability in the precise locations of functional areas, and ii) the proximity of language and MD networks in the frontal and parietal cortex. These factors would lead to the blurring of language/MD network boundaries, critically undermining our ability to separate these networks in the analyses (see e.g., Nieto-Castanon & Fedorenko, 2012, and Fedorenko & Blank, 2020, for discussion). That said, as noted below, we make the individual contrast maps publicly available, allowing others to re-analyze the current dataset using any additional approaches, to complement the set of analyses reported here.)

### Participants

Four hundred and eighty-one unique individuals (age 18-71, mean 26.4; 288 (∼60%) females; see **Table SI-1** available at OSF: https://osf.io/pdtk9/ for information about participants’ age, sex, and handedness) from the Cambridge/Boston, MA community participated for payment across 30 fMRI language comprehension experiments, with 11-385 participants per experiment (see **Table 1** for numbers of participants in each experiment; see **Table SI-4** at https://osf.io/pdtk9/ for information about participant overlap among experiments). Each participant completed 1-14 critical experiments (median=1), for a total of 678 critical experiment scanning sessions comprising the current dataset (see below for details). Four hundred and fifty-five participants (∼95%) were right-handed, as determined by the Edinburgh handedness inventory (Oldfield, 1971), or self-report; the remaining 26 left-handed participants showed typical left-lateralized language activations in the language localizer task (see Willems et al., 2014, for arguments for including left-handers in cognitive neuroscience experiments). Four hundred and two participants (∼83%) were native speakers of English; the remaining 79 participants were native speakers of diverse languages and fluent speakers of English (for these participants, we examined responses to language processing in their native language; data from Ayyash, Malik-Moraleda, et al., in prep.). All participants gave informed consent in accordance with the requirements of MIT’s Committee on the Use of Humans as Experimental Subjects (COUHES).

**Table 1.**
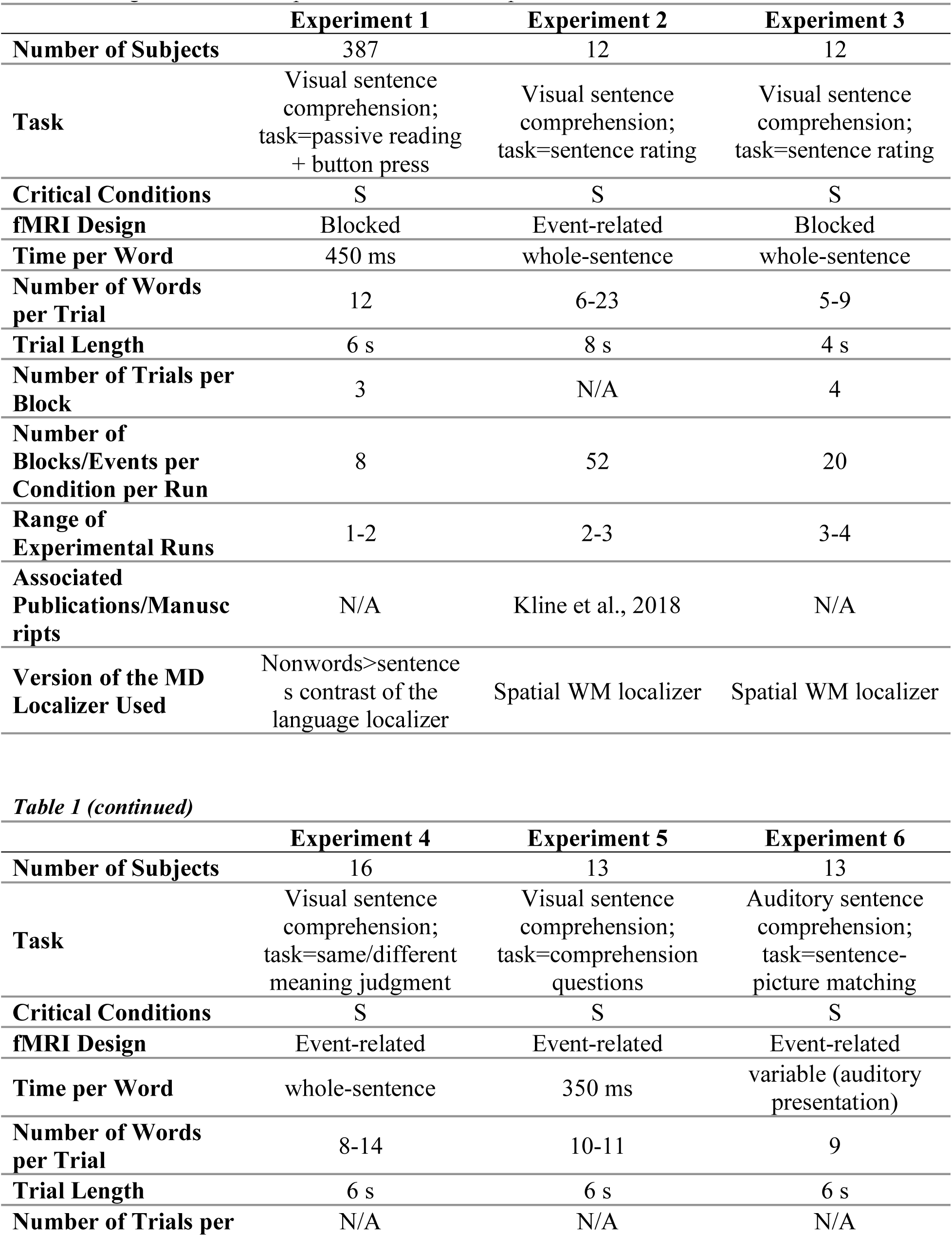
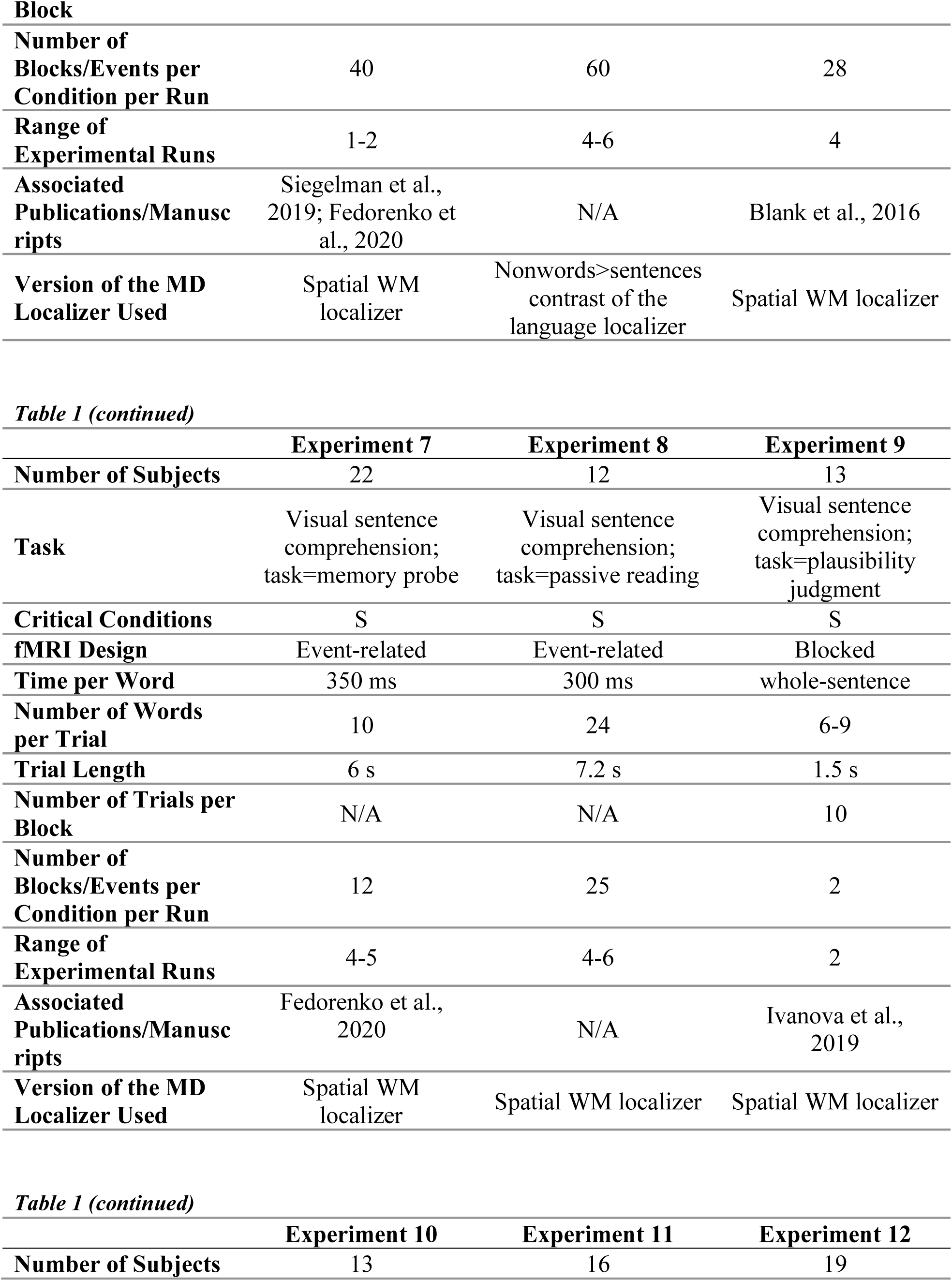
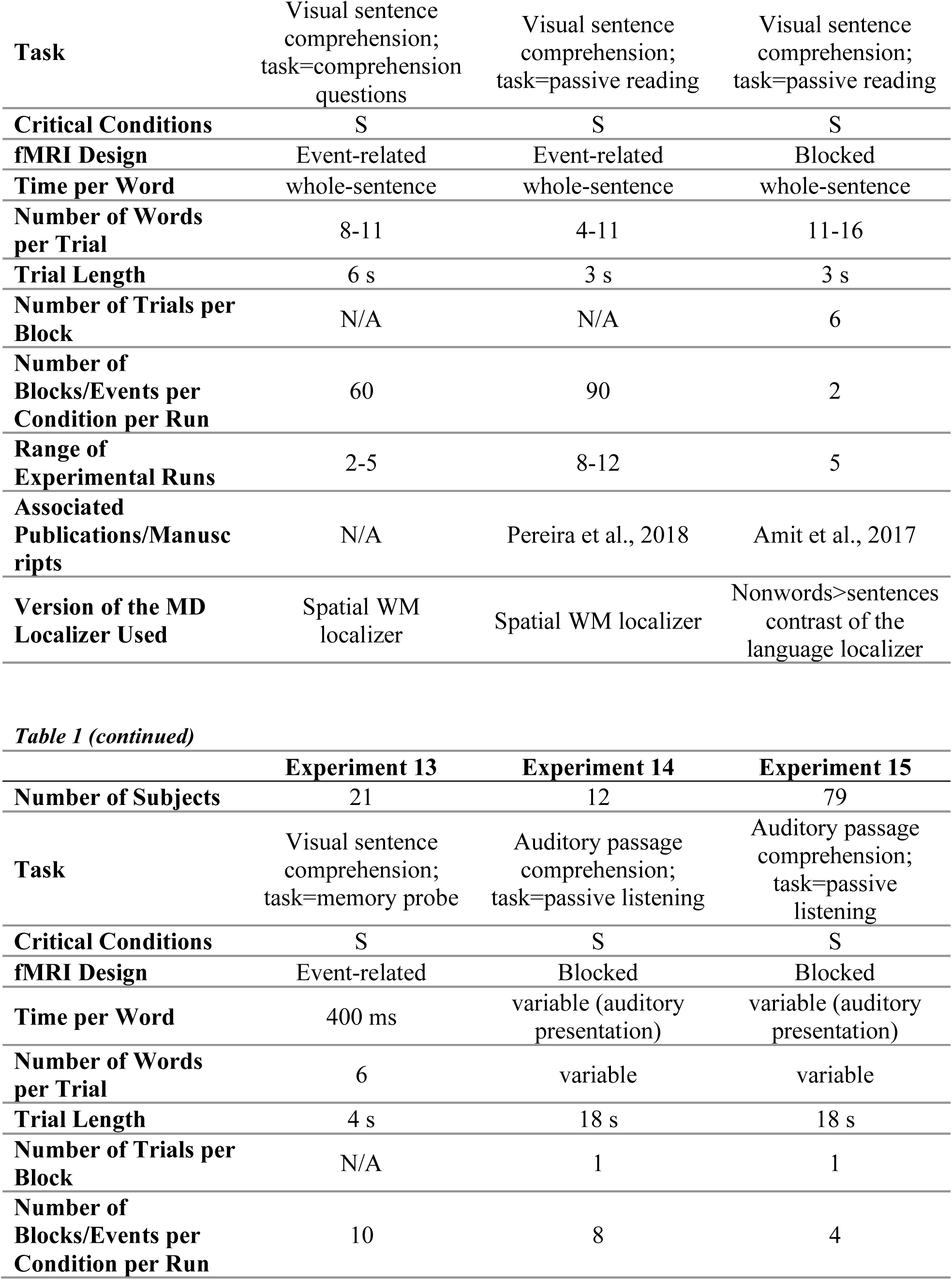
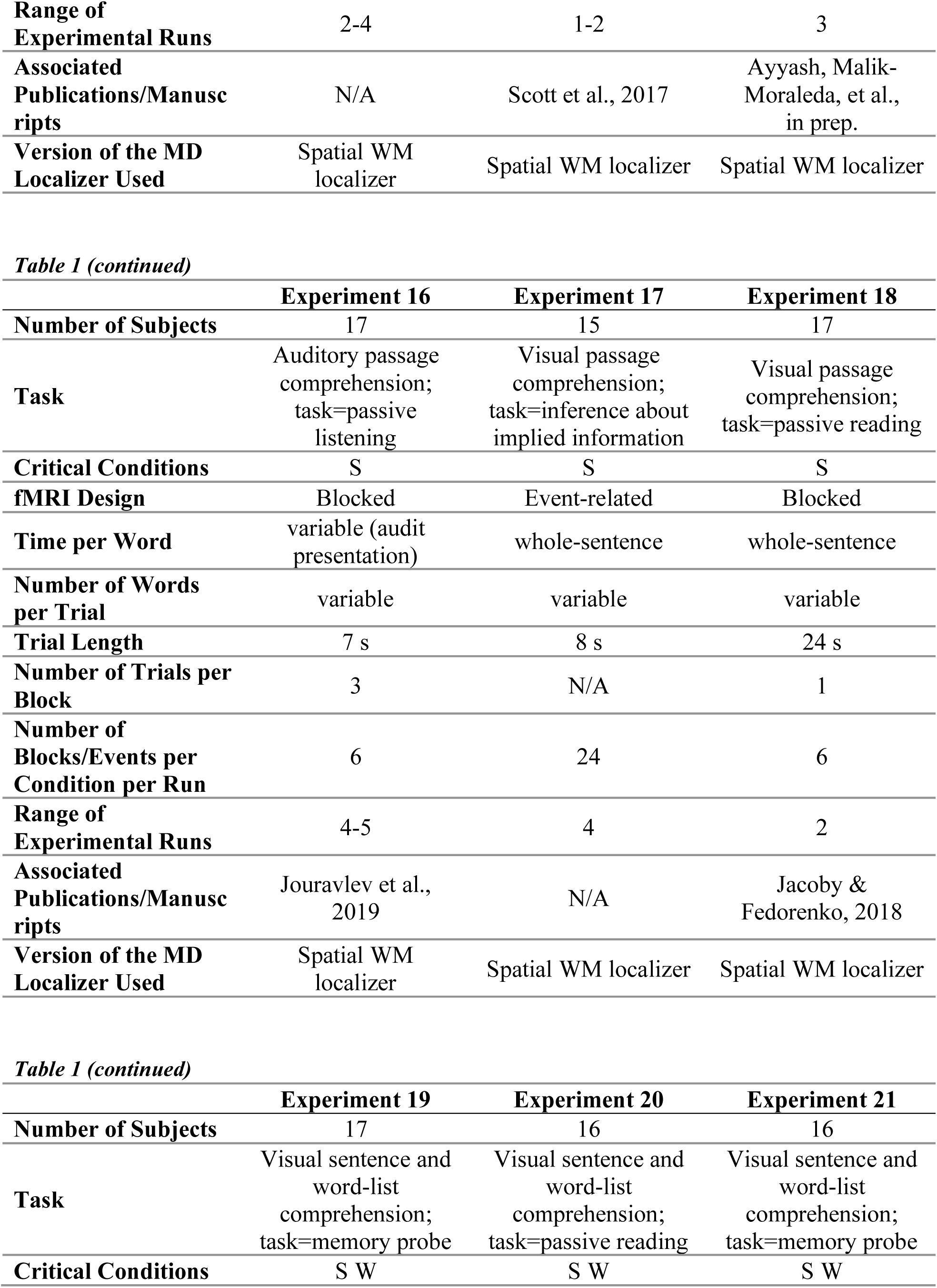
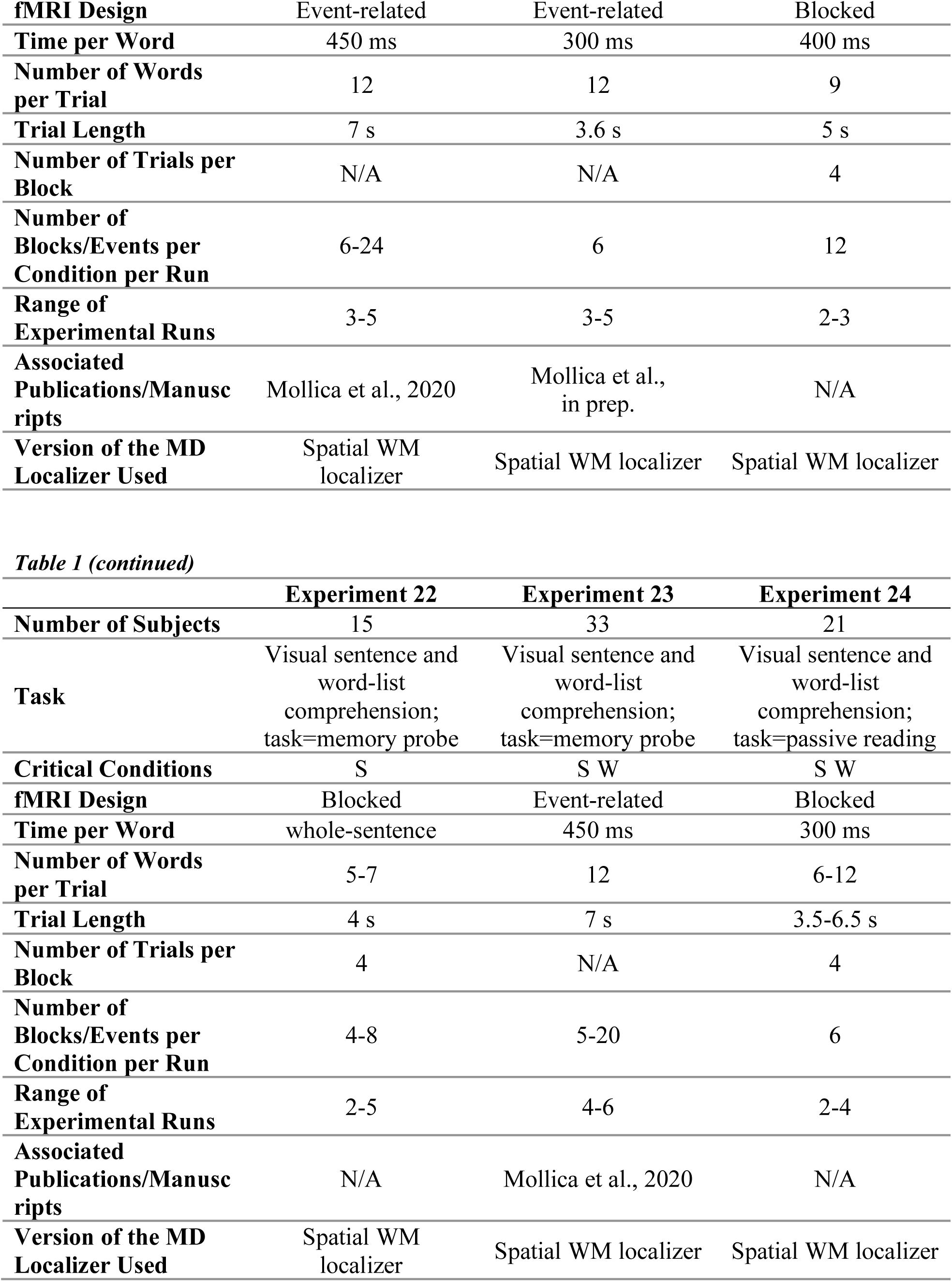
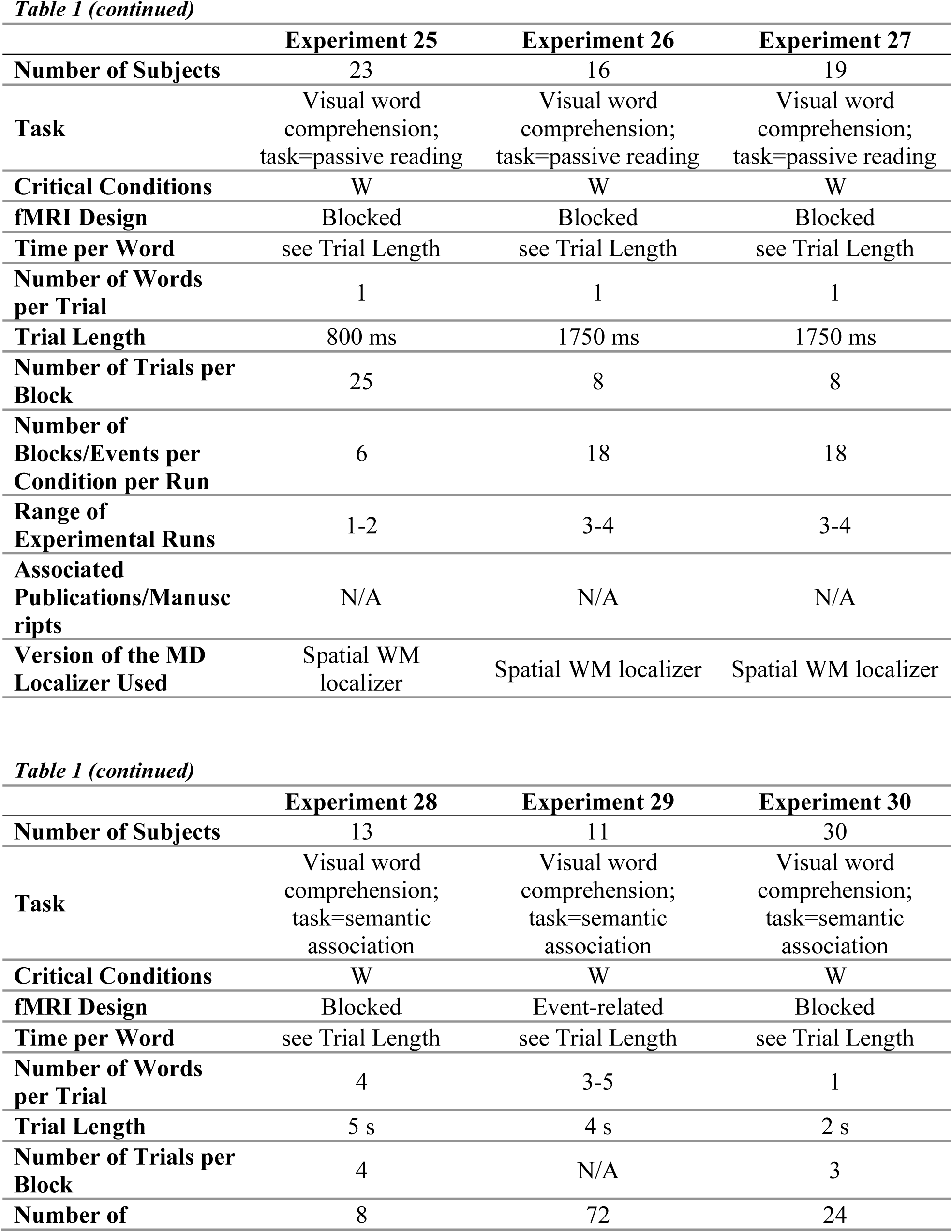
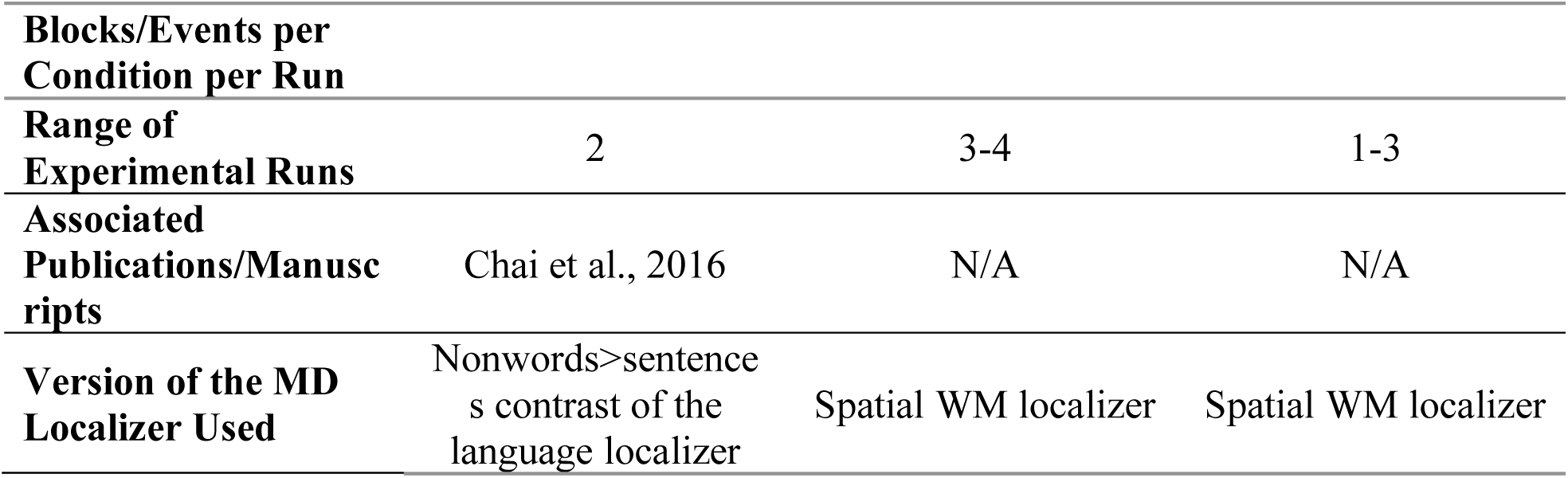
Design, materials, and procedure details for Experiments 1-30

### Design, stimuli, and procedure

In describing the dataset in more detail, it is helpful to define a few terms. A ***critical experiment dataset*** is a set of functional runs for a single participant for a critical experiment (total number of critical experiments = 30). A ***(scanning) session*** is a single visit of a participant to the MRI facility, during which one or more experiments are run. A ***critical experiment session*** is a session that contains one or more critical experiment datasets. An ***MD localizer session*** is a session that contains data for an MD localizer (one of two versions, as detailed below). A ***language localizer session*** is a session that contains data for a language localizer.

We have 939 critical experiment datasets (see **Table 1**) across 678 critical experiment sessions. For 26 of the 30 experiments (507/939 critical experiment datasets), we functionally identified the MD network using a spatial working memory (WM) localizer described below (e.g., Blank et al., 2014). For the remaining 4 experiments (432/939 critical experiment datasets), we used another difficulty manipulation based on a contrast between the reading of nonwords and the reading of sentences, as in Fedorenko et al. (2013). Furthermore, for the 26 experiments that used the spatial WM MD localizer, in 307 of the 507 critical experiment datasets the MD localizer was administered in the same scanning session as the critical experiment; in the remaining 200 critical experiment datasets, the MD localizer came from an earlier scanning session (the activation patterns are highly stable within and across scanning sessions; Assem et al., 2019, Shashidhara et al., 2020, unpublished data from the Fedorenko lab). Similarly, for the 4 experiments that used the *nonwords > sentences* MD localizer contrast, in 418 of the 432 critical experiment datasets the MD localizer was administered in the same scanning session as the critical experiment; in the remaining 14 critical experiment datasets, the MD localizer came from an earlier scanning session.

All participants further completed a language localizer task (Fedorenko et al., 2010). The language functional regions of interest (fROIs) were used in some control analyses, as detailed below. One version of the language localizer served as one of the critical language experiments given that it included a passive sentence comprehension condition. In 748/939 critical experiment datasets, the language localizer was administered in the same scanning session as the critical experiment; in the remaining 191 critical experiment datasets, the language localizer came from an earlier scanning session (the activation patterns are highly stable within and across scanning sessions; Mahowald & Fedorenko, 2016).

Most scanning sessions lasted approximately 2 hours and included one or more other tasks for unrelated studies.

#### MD localizer

For 26/30 critical experiments (507/939 critical experiment datasets), regions of the MD network were localized using a spatial working memory (WM) task contrasting a harder condition with an easier condition (e.g., Fedorenko et al., 2011, 2013; Blank et al., 2014). On each trial (8 s), participants saw a fixation cross for 500 ms, followed by a 3×4 grid within which randomly generated locations were sequentially flashed (1 s per flash) two at a time for a total of eight locations (hard condition) or one at a time for a total of four locations (easy condition). Then, participants indicated their memory for these locations in a two-alternative, forced-choice paradigm via a button press (the choices were presented for 1,000 ms, and participants had up to 3 s to respond). Feedback, in the form of a green checkmark (correct responses) or a red cross (incorrect responses), was provided for 250 ms, with fixation presented for the remainder of the trial. Hard and easy conditions were presented in a standard blocked design (4 trials in a 32 s block, 6 blocks per condition per run) with a counterbalanced order across runs. Each run included 4 blocks of fixation (16 s each) and lasted a total of 448 s (**Figure 1**). The *hard > easy* contrast targets brain regions engaged in cognitively demanding tasks. Fedorenko et al. (2013) have established that the regions activated by this task are also activated by a wide range of other demanding tasks (see also Duncan & Owen, 2000; Hugdahl et al., 2015). For the remaining 4 critical experiments in which not every participant performed the spatial WM task (432/939 of the critical experiment datasets), we used the *nonwords > sentences* contrast of the language localizer task (described below) to define the MD fROIs (Fedorenko et al., 2013).

**Figure 1.**
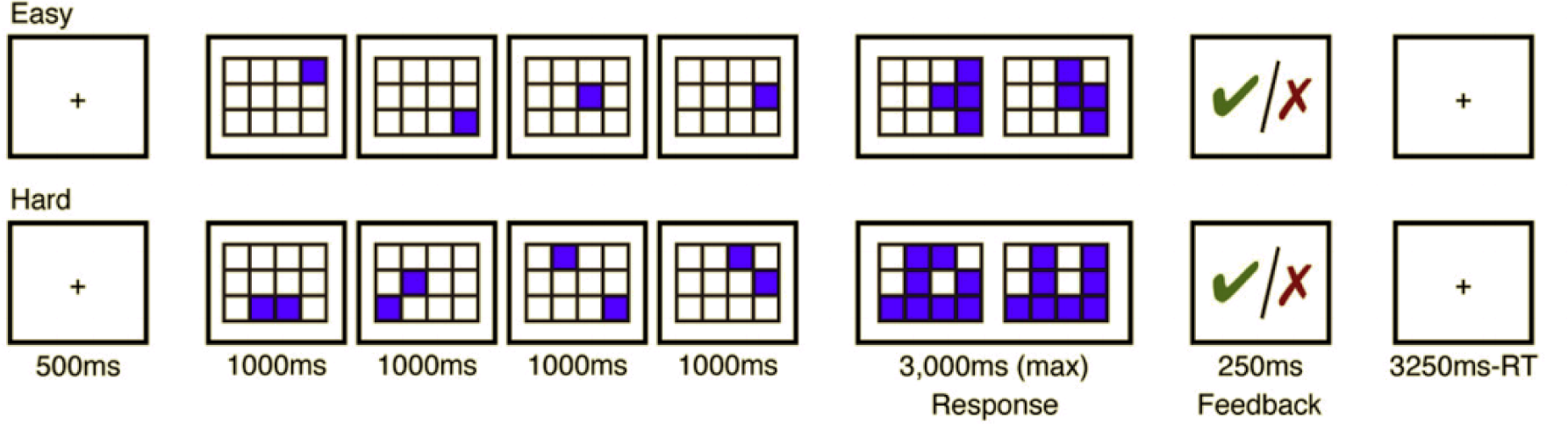
Procedure and timing for the spatial working memory task used to localize the multiple demand (MD) fROIs.

#### *Language localizer* (used in some control analyses, and as one of the critical experiments)

This task is described in detail in Fedorenko et al. (2010). Briefly, participants read sentences and lists of unconnected, pronounceable nonwords in a blocked design. Stimuli were presented one word/nonword at a time. Each of the 481 unique participants completed one or more language localizer sessions (n=423 completed a single localizer session; n=46 completed 2 sessions; n=8 completed 3 sessions; and n=4 completed 4 sessions), for a total of 555 language localizer sessions included in the analyses. Across this dataset, five slightly different versions of the language localizer were used (see **Table 2**). For 71 language localizer sessions, each trial ended with a memory probe and participants had to indicate, via a button press, whether or not that probe had appeared in the preceding sentence / nonword sequence. In the remaining 484 localizer sessions, participants read the materials passively and performed a simple button-press task at the end of each trial (included in order to help participants remain alert). The language localizer has been shown to be robust to changes in the materials, modality of presentation, and task (Fedorenko et al., 2010; Fedorenko, 2014; Scott et al., 2017; Ivanova, Siegelman, et al., in prep.).

**Table 2.**
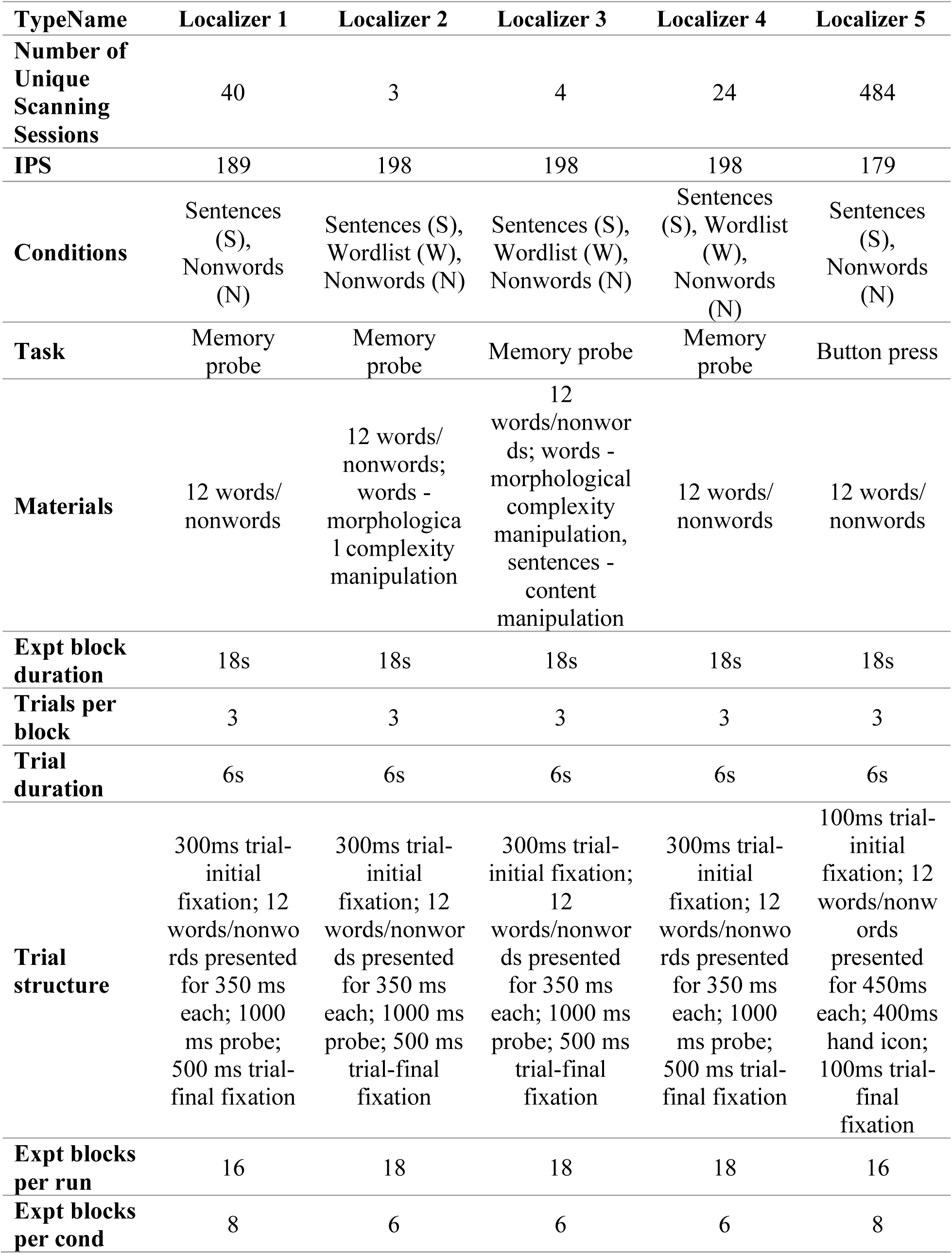
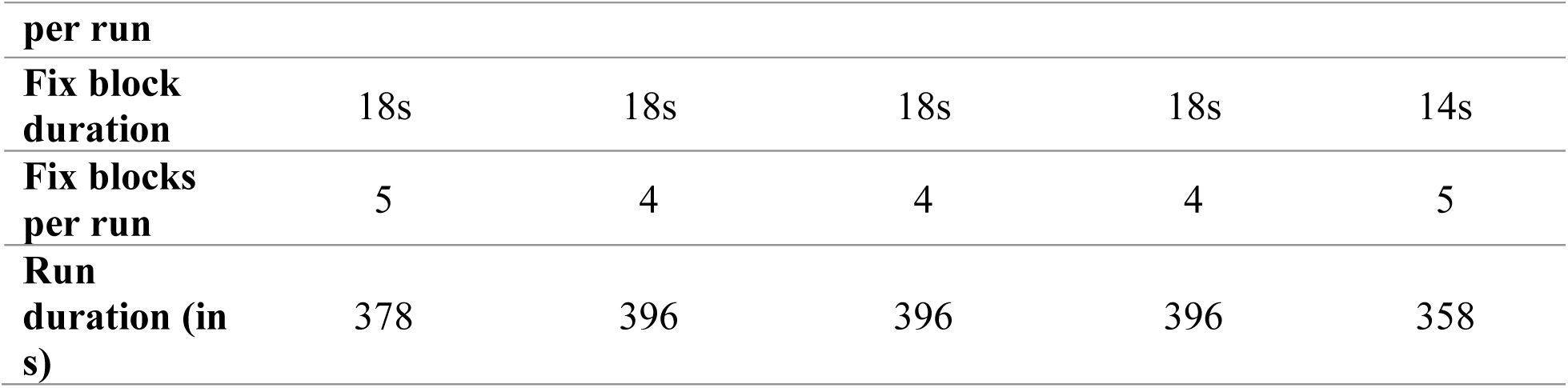
Timing parameters for the different versions of the language localizer task.

#### Critical experiments

To broadly evaluate the role of the MD network in language comprehension, we examined neural responses across 30 diverse experiments conducted in the Fedorenko lab between 2010 and 2019, which included word-level and sentence-level materials. Details of all the experiments are reported in **Table 1**, but we here summarize the general approach to the selection of experimental conditions and the key dimensions of variation present across the experiments.

Each of the 30 experiments was originally designed to evaluate a specific hypothesis about (i) the sensitivity of the language and/or the MD network to some linguistic (lexical, syntactic, semantic, or pragmatic) manipulation, or (ii) the selectivity of the two networks for linguistic vs. non-linguistic conditions. For example, Experiment 2 compared responses to one-liner jokes vs. closely matched non-joke controls (Kline et al., submitted); Experiment 6 compared responses to syntactically simper vs. more complex sentences (sentences containing subject-vs. object-extracted relative clauses) (Blank et al., 2016); Experiment 16 compared responses to spoken linguistic materials vs. speech-accompanying gestures (Jouravlev et al., 2019); and Experiment 24 contrasted sentences that contained a temporary syntactic ambiguity vs. control unambiguous sentences (following the design of Snijders et al., 2009). Data from some of these experiments have been published or are reported in preprints and papers under review (see **Table 1**); other experiments are parts of ongoing projects and have not yet been reported anywhere (we make the data used in the analyses below—estimates of neural responses to the conditions of the critical and localizer experiments in the MD and language fROIs in each participant (**Table SI-2**)—as well as whole-brain activation maps for the critical and localizer contrasts, available at https://osf.io/pdtk9/; raw data are available from the senior author upon request). For the purposes of this study, in each experiment, we (i) selected only the conditions where participants were asked to read or listen to words / word lists or plausible well-formed sentences (we excluded conditions that e.g., contained syntactic violations), and, where necessary, (ii) averaged the responses across the fine-grained linguistic manipulations to derive a single response magnitude for (a) word comprehension and/or (b) sentence comprehension.

Eighteen experiments involved sentence comprehension (3 of these involved passages, 14 – unconnected sentences, and 1 – both passages and unconnected sentences), six involved word-level comprehension, and the remaining six contained both sentence materials and matched word-list conditions. In 26 experiments, linguistic materials were presented visually, and in the remaining 4 – auditorily. Critically, for the research question asked here, the experiments varied in the task used: in 13 experiments, participants read or listened to the materials passively (sometimes accompanied by a simple button-press task), and in the remaining 17 experiments, they were asked to perform a task (a memory probe task in 6 experiments, a semantic association task in 3 experiments, a sentence rating task in 2 experiments, a comprehension-question task in 2 experiments, a meaning similarity judgment task in 1 experiment, an inference task in 1 experiment, a plausibility judgment task in 1 experiment, and a sentence-picture matching task in 1 experiment).

To summarize some of the procedural/timing details (provided in **Table 1**), 16 experiments used a blocked design, and the other 14 – an event-related design. In blocked design experiments, participants saw or heard between 4 and 72 blocks per condition (each between 8.5 and 26 s in duration). (Note that “condition” here is the overarching sentence-comprehension or word-comprehension condition; so, for example, if an experiment had two conditions – syntactically easy and syntactically more complex sentences – we here report the number of blocks across the two conditions, given that we average the responses between those two conditions in the analyses, as described above.) In event-related design experiments, participants saw or heard between 18 and 1,080 trials per condition (each between 3 and 8 s in duration). The materials for all experiments are available from the senior author upon request (those that come from published studies are typically available on the associated OSF pages, as indicated in the relevant publications).

### Data acquisition, preprocessing, and first-level modeling

#### Data acquisition

Whole-brain structural and functional data were collected on a whole-body 3 Tesla Siemens Trio scanner with a 32-channel head coil at the Athinoula A. Martinos Imaging Center at the McGovern Institute for Brain Research at MIT. T1-weighted structural images were collected in 176 axial slices with 1 mm isotropic voxels (repetition time (TR) = 2,530 ms; echo time (TE) = 3.48 ms). Functional, blood oxygenation level-dependent (BOLD) data were acquired using an EPI sequence with a 90° flip angle and using GRAPPA with an acceleration factor of 2; the following parameters were used: thirty-one 4.4 mm thick near-axial slices acquired in an interleaved order (with 10% distance factor), with an in-plane resolution of 2.1 mm × 2.1 mm, FoV in the phase encoding (A >> P) direction 200 mm and matrix size 96 × 96 voxels, TR = 2,000 ms and TE = 30 ms. The first 10 s of each run were excluded to allow for steady state magnetization.

#### Preprocessing

Data preprocessing was carried out with SPM5 (using default parameters, unless specified otherwise) and supporting, custom MATLAB scripts (available from the senior author upon request). (We used an older version of SPM here because the current dataset is part of a larger dataset with ∼900 participants and over 2,000 scanning sessions, and we wanted to keep the preprocessing and first-level modeling consistent across that larger dataset; but see below.) Preprocessing of anatomical data included normalization into a common space (Montreal Neurological Institute (MNI) template), resampling into 2 mm isotropic voxels. Preprocessing of functional data included motion correction (realignment to the mean image using 2^nd^-degree b-spline interpolation), normalization (estimated for the mean image using trilinear interpolation), resampling into 2 mm isotropic voxels, smoothing with a 4 mm FWHM Gaussian filter and high-pass filtering at 200 s.

#### First-level modeling

For both the MD localizer task and the critical tasks, a standard mass univariate analysis was performed in SPM5, separately for each participant, whereby a general linear model (GLM) estimated, for each voxel, the effect size of each condition in each experimental run. These effects were each modeled with a boxcar function (representing entire blocks/events) convolved with the canonical Hemodynamic Response Function (HRF). The model also included first-order temporal derivatives of these effects, as well as nuisance regressors representing entire experimental runs and offline-estimated motion parameters.

To ensure that the results are robust to the version of SPM used for preprocessing and first-level modeling, we re-analyzed four of the thirty experiments in SPM12. Although the mean responses in the MD fROIs to the critical conditions differed slightly between the two pipelines, they i) were overall very similar, and ii) did not exhibit any systematic bias, which could have affected any of the critical comparisons (see **Figure SI-4** at https://osf.io/pdtk9/, showing a direct pipeline comparison for these experiments).

### Definition of the MD functional regions of interest (fROIs)

For each critical experiment dataset of each participant, we defined a set of multiple demand (MD) functional ROIs using group-constrained, subject-specific localization (Fedorenko et al., 2010). In particular, as described above, for 26 of the 30 experiments (507/939 critical experiment datasets), we used the spatial working memory (WM) MD localizer. Each individual map for the *hard > easy* spatial WM contrast was intersected with a set of twenty binary masks. These masks (**Figure 2**; masks available for download from https://osf.io/pdtk9/) were derived from a probabilistic activation overlap map for the same contrast in a large set of participants (n=197) using watershed parcellation, as described in Fedorenko et al. (2010), and corresponded to relatively large areas within which most participants showed activity for the target contrast. These masks covered the fronto-parietal MD network (including what some treat as a separate, cingulo-opercular, sub-network; e.g., Power et al., 2011), and were highly overlapping with a set of anatomical masks used in Fedorenko et al. (2013). For the remaining 4 experiments (432/939 critical experiment datasets), we used a contrast between the reading of nonwords and the reading of sentences. Each individual map for the *nonwords > sentences* contrast was intersected with the same twenty masks (see Fedorenko et al., 2013, for evidence that this contrast yields similar activations to more typical executive function tasks). Within each mask, a participant-specific MD fROI was defined as the top 10% of voxels with the highest *t*-values for the localizer contrast (*hard > easy* or *nonwords > sentences*).

**Figure 2.**
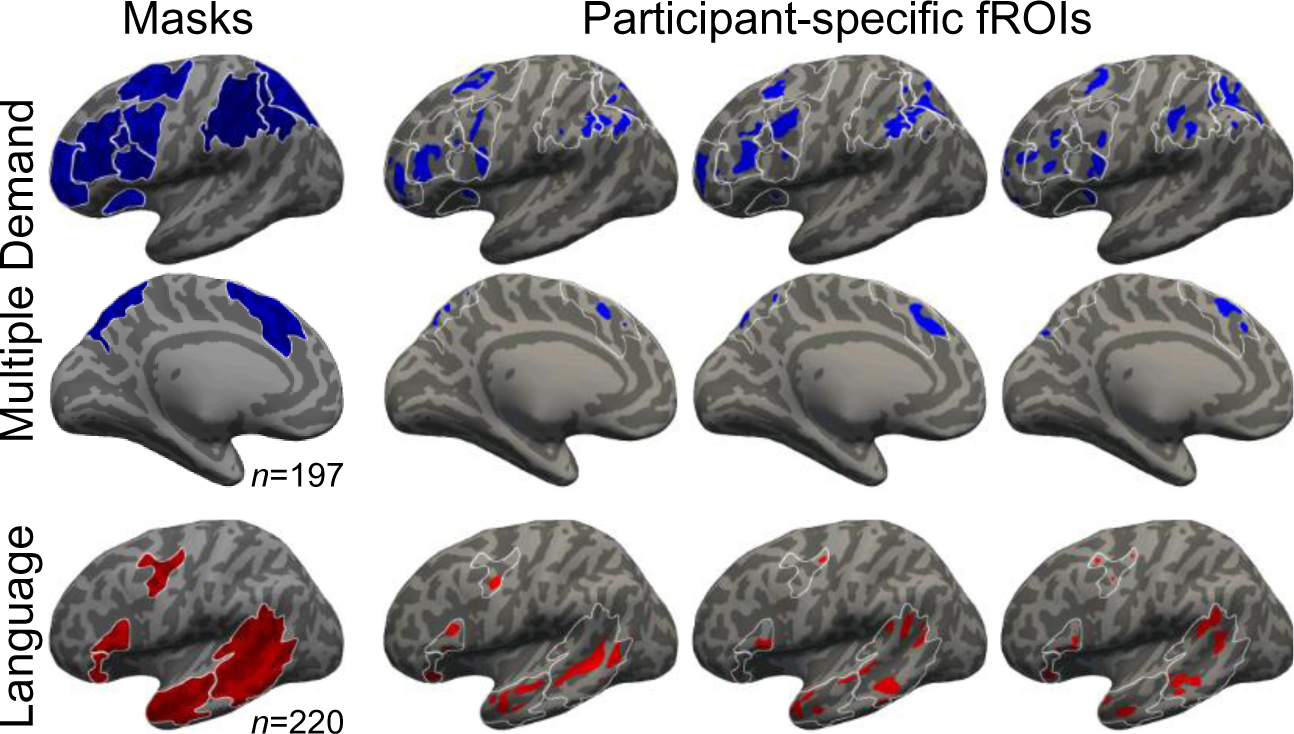
Masks and subject-specific functional regions of interest (fROIs). Data are shown for the multiple-demand (MD) network (top, middle; blue) and language network (bottom; red), and are approximate projections from functional volumes onto the cortical surface of an inflated average brain in common space. Only the left hemisphere is shown. The leftmost column shows masks derived from a group-level representation of data for the MD localizer contrast (Hard>Easy) and the language localizer contrast (Sentences >Nonwords), in an independent group of subjects, using watershed parcellation. These masks were used to constrain the selection of subject-specific fROIs. The other columns show approximate locations example of MD and language fROIs from 3 subjects. Apparent overlap across MD and language fROIs within an individual is illusory and due to projection onto the cortical surface. Note that, because data were analyzed in volume (not surface) form, some parts of a given fROI that appear discontinuous in the figure (e.g., separated by a sulcus) are contiguous in volumetric space. White contours denote the borders of the masks.

In all the critical analyses reported here, we treat the MD fROIs as a functionally integrated system given that prior work has established that these regions not only share functional profiles, but also that the MD regions’ time-courses are strongly correlated during both rest and task performance (e.g., Blank et al., 2014; Paunov et al., 2019), and the effect sizes in task-based paradigms correlate strongly across participants (Mineroff et al., 2018; Assem et al., 2019). However, we acknowledge the possibility that subdivisions may exist within this network (e.g., Blank et al., 2014; Paunov et al., 2019). And treating this network as an integrated system need not imply that all of its regions are identical in their response patterns and functions.

#### Definition of the language (fROIs) (for control analyses)

To define the language fROIs, each individual map for the *sentences > nonwords* contrast from the language localizer was intersected with a set of five binary masks. These masks (**Figure 2**; masks available for download from https://osf.io/pdtk9/) were derived from a probabilistic activation overlap map for the language localizer contrast in a large set of participants (n=220), following the method described in Fedorenko et al. (2010) for a smaller set of participants. These masks covered the fronto-temporal language network in the left hemisphere.

### Validation of the MD fROIs

To ensure that the MD fROIs behave as expected (i.e., show a reliably greater response to the hard spatial WM condition compared to the easy one, or a greater response to the nonwords condition compared to the sentences condition), we used an across-runs cross-validation procedure (e.g., Nieto-Castañon & Fedorenko, 2012). For the 26 experiments in which all participants completed the spatial WM MD localizer task, we identified the unique participants that completed two runs of the MD localizer, leaving us with 172 sessions for the cross-validation analysis. Similarly, for the 4 experiments where we used the *nonwords > sentences* MD localizer task, we identified the unique participants that completed two runs of the MD localizer, leaving us with 366 sessions for the cross-validation analysis. In this analysis, the first run of the localizer was used to define the fROIs, and the second run to estimate the responses (in percent BOLD signal change, PSC) to the localizer conditions, ensuring independence (e.g., Kriegeskorte et al., 2009); then the second run was used to define the fROIs, and the first run to estimate the responses; finally, the extracted magnitudes were averaged across the two runs to derive a single response magnitude for each of the localizer conditions (see **Table SI-3a** at https://osf.io/pdtk9/; the for the cross-validated responses to the language localizer conditions in the language fROIs, see Table SI-3b). Statistical analyses were performed on these extracted PSC values (the MATLAB code is available at https://osf.io/pdtk9/). For the 26 MD localizer sessions that only contained a single run of the spatial WM task, we used visual examination of whole-brain activation maps for the *hard > easy* contrast, to ensure that the expected pattern of activation is observed.

### Critical analyses

To estimate the responses in the MD fROIs to the conditions of the critical experiments, the data from all the runs of the MD localizer were used to define the fROIs, and the responses to each condition (sentence comprehension and/or word comprehension) were then estimated in these regions, and, in some cases, averaged across conditions to derive a single response magnitude for sentence comprehension and/or word comprehension, as described above. Statistical analyses were then performed on these extracted PSC values (see **Table SI-2** and the associated R code at https://osf.io/pdtk9/).

To characterize the role of the MD network in language comprehension, we ran several linear mixed effect models using the “lme4” package in R with *p*-value approximation performed by the “lmerTest” package (Bates et al., 2015; Kuznetsova et al., 2017). In particular, we asked four questions. First, we asked whether—across experiments—the MD network is engaged (above the low-level fixation baseline) during language comprehension. Second, we asked whether the MD network, like the language network, shows stronger responses to language processing in the left compared to the right hemisphere. Third, we compared the MD network’s responses to sentences vs. words / word lists. One robust signature of the language network is a stronger response to sentences compared to word lists (e.g., Snijders et al., 2009; Fedorenko et al., 2010; Pallier et al., 2011; Fedorenko et al., 2016), presumably because processing sentences requires additional computations compared to processing individual word meanings (see e.g., Mollica et al., 2020, for a recent discussion). We wanted to test whether the MD network shows a similar preference for sentences. Finally and critically, we asked whether the MD network’s engagement is stronger for experiments that had included an explicit task, compared to the ones where participants passively read or listened to stimuli.

### *\** Is the MD network engaged during language comprehension?

*Effect size ∼ condition + (1+condition|ID) + (1+condition|ROI) + (1+condition|experiment)*

We fit a linear mixed effect regression model, predicting the level of BOLD response in the MD fROIs across the thirty experiments. The model included a fixed effect for condition (sentences vs. words / word lists; the difference between these levels was not of interest to this particular question, and was included for appropriately modeling variance in the data). In addition, it included random intercepts and slopes for condition by participant, fROI, and experiment.

### *\** Does the MD network show left-lateralized responses?

*Effect size ∼ hemisphere + (1+hemisphere|ID) + (1+hemisphere|ROI) + (1+hemisphere|experiment)*

We fit a linear mixed effect regression model, predicting the level of BOLD response in the MD fROIs across experiments, separately for sentence conditions and word conditions. (We tested the model for sentence and word conditions separately because the hemisphere×condition interaction was significant in a combined model.) The model included a fixed effect for hemisphere and random intercepts and slopes for hemisphere by participant, fROI, and experiment. The mean difference between the fixed effects for the two hemispheres was tested against zero using the “glht” command in “multcomp” package (Hothorn et al., 2008) in R.

Additionally, we performed the same analysis for the language network fROIs, which are expected to show robust left lateralization (e.g., Mahowald & Fedorenko, 2016).

It is worth noting that directly comparing parameter estimates from the homologous regions in the two hemispheres is potentially problematic given the inter-hemispheric differences in vascularization and hemodynamic response properties (e.g., Handwerker et al., 2004; Miezin et al., 2010; Hedna et al., 2013; cf. Taylor et al., 2018). In particular, the use of a canonical HRF in modeling neural responses (albeit with time derivatives) may affect the degree of model fit in the two hemispheres, which would contribute to differences in the strengths of response, thus complicating interpretation. However, because no clear alternatives exist for comparing activity between the hemispheres, we chose to include these analyses.

### *\** Does the MD network respond differentially to sentences vs. words / word lists?

*Effect size ∼ condition + (1+condition|ID) + (1+condition|ROI) + (1+condition|experiment)*

We fit a linear mixed effect regression model, predicting the level of BOLD response in the MD fROIs across the thirty experiments. The model included a fixed effect for condition (sentences vs. words / word lists), and random intercepts and slopes for condition by participant, fROI, and experiment. The mean difference between the fixed effects for sentences vs. words / word lists was tested against zero using the “glht” command in “multcomp” package (Hothorn et al., 2008) in R.

Additionally, we performed the same analysis for the language network fROIs, which are expected to show a robust *sentences > word lists* effect (e.g., Snijders et al., 2009; Fedorenko et al., 2010; Pallier et al., 2010; Fedorenko et al., 2016).

### *\** Does the MD network respond differentially to language comprehension depending on whether an explicit task is used?

*Effect size ∼ task + (1|ID) + (1+task|ROI) + (1|experiment)*

We fit a linear mixed effect regression model, predicting the level of BOLD response in the MD fROIs across the thirty experiments. The model included a fixed effect for the type of task that participants had to perform (passive reading/listening vs. an active task), random intercepts by participant and experiment, as well as a random intercept and slope for task by fROI. The mean difference between the fixed effects for active vs. passive task was tested against zero using the “glht” command in “multcomp” package (Hothorn et al., 2008) in R.

Additionally, we performed the same analysis for the language network fROIs. Whether/how language regions are modulated by the presence of a task is debated (e.g., Roskies et al., 2001; Noesselt et al., 2003; Andoh & Paus, 2011), so we took an opportunity to use this rich dataset to shed light on this question.

## Results

### Validation of the MD fROIs

As expected, and replicating prior work (e.g., Fedorenko et al., 2013; Blank et al., 2014; Mineroff et al., 2018; Assem et al., 2019), each of the MD fROIs showed a highly robust *hard > easy* effect (all *t*s(171)>18.7; *p*s<10^-72^, FDR-corrected for the twenty ROIs; Cohen *d*s > 0.65, based on a dependent-samples *t*-test). Similarly, for the participants for whom the *nonwords > sentences* contrast was used to define the MD fROIs, each of the fROIs showed a robust *nonwords > sentences* effect (*t*s(365)>12; *p*s<10^-68^, FDR-corrected for the twenty ROIs; Cohen *d*s > 0.41, based on a dependent-samples *t*-test).

### Critical results

Replicating numerous prior studies that have reported activation within the MD network for linguistic manipulations (e.g., Kuperberg et al., 2003; Rodd et al., 2005; Novais-Santos et al., 2007; January et al., 2009; Peelle et al., 2010; Nieuwland et al., 2012; McMillan et al., 2013; Mollica et al., 2020), we found that—across experiments— language comprehension tasks elicited an above baseline response in the MD network (**Figure 3**) (sentences: b = 0.27, SE = 0.09, z = 2.91, p = 0.003; words / word lists: b = 0.41, SE = 0.07, z = 6.32, p < 10^-9^). Additionally, we found that the MD fROIs in the left hemisphere responded more strongly than the MD fROIs in the right hemisphere, for both sentence and word-level comprehension (**Figure 3**) (sentences: b = 0.20, SE = 0.04, z = 4.72, p < 10^-5^; words: b = 0.18, SE = 0.06, z = 2.98, p = 0.003). As expected, this pattern was also robustly present in the language network (sentences: b = 0.62, SE = 0.14, z = 4.45, p < 10^-5^; words: b = 0.40, SE = 0.09, z = 4.23, p < 10^-4^).

**Figure 3.**
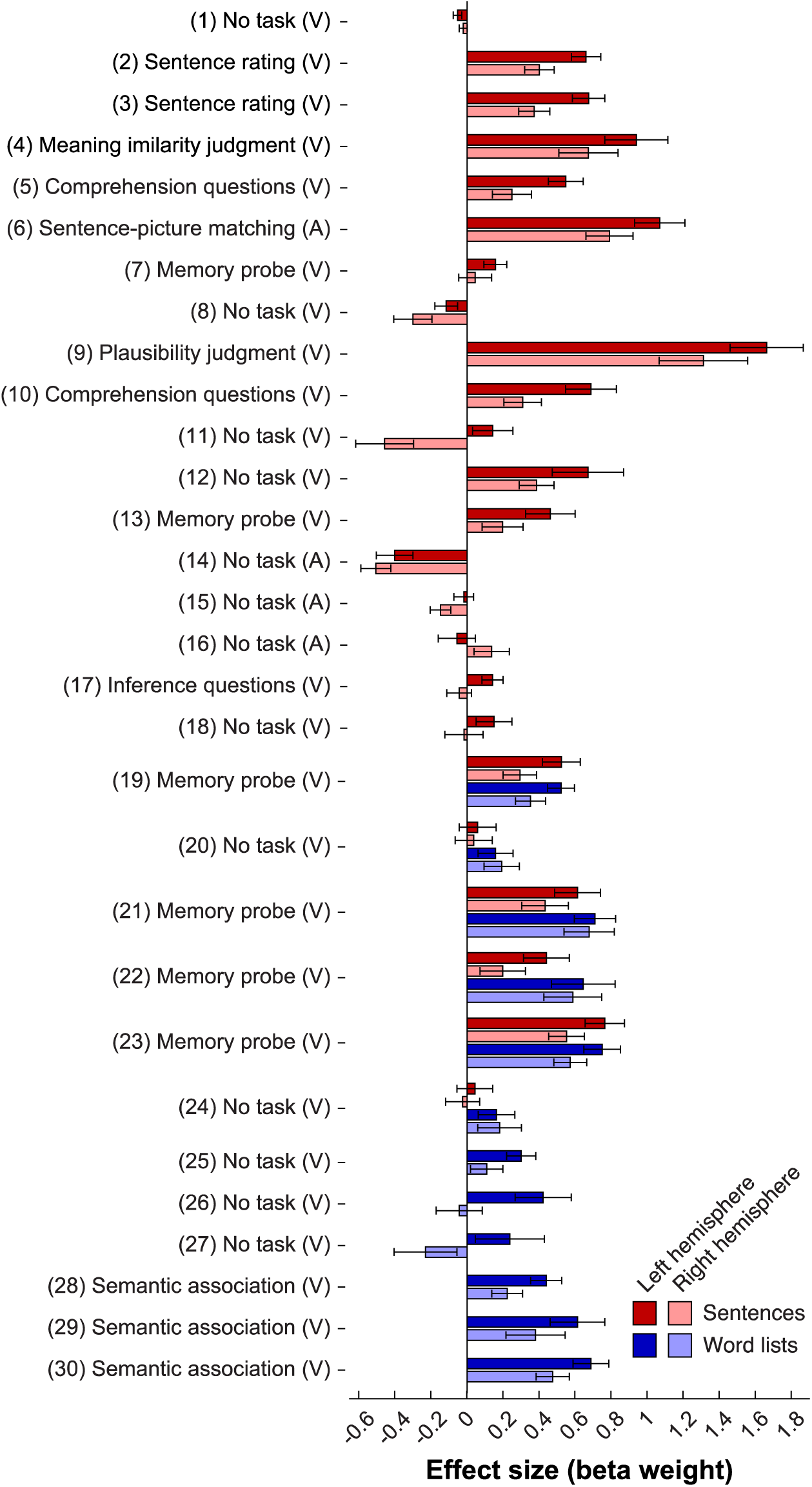
Responses to sentences and word lists in the MD network across experiments, and their laterality. Responses (beta weights for the corresponding regressors in the GLM) are shown averaged across fROIs in the left (darker colors) and right (brighter colors) hemispheres, separately for sentences (red shades) and word lists (blue shades) conditions. Responses are measured as beta weights for the corresponding condition regressors in the GLM. Bars show the average response across subjects, and errors bars show standard errors of the mean across subjects. Most experiments include either sentences or word lists, except for experiments 19-24.

However, in contrast to the language network, which responds more strongly during sentence comprehension compared to the processing of unconnected lists of words (e.g., Fedorenko et al., 2010), an effect we replicated here (b = 0.37, SE = 0.10, z = 3.51, p = 0.0004), the MD network showed the opposite pattern, with a stronger response to words / word lists than sentences (b = 0.15, SE = 0.06, z = 2.55, p = 0.011; **Figure 4**; see **Figure SI-1** for responses of individual fROIs).

**Figure 4.**
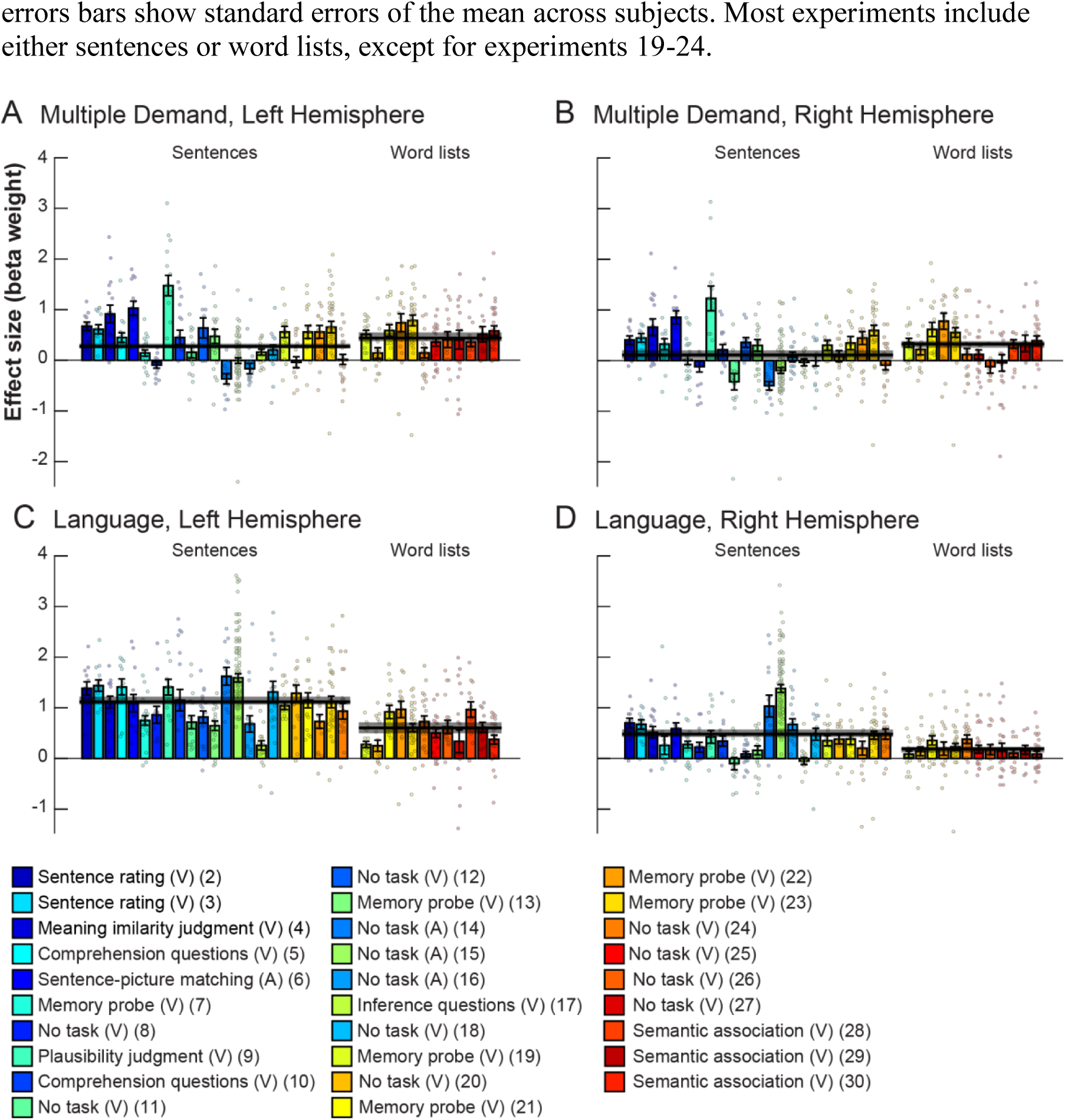
Responses of each network and hemisphere to the sentences and words / word lists conditions across experiments. Responses (beta weights for the corresponding regressors in the GLM) are shown averaged across fROIs in the MD (top) and language (bottom) networks, separately for the left hemisphere (left) and right hemisphere (right). Data are presented for each of 29 experiments (data for Experiment 1 are not shown, because the number of individual data points was too large for the plot to be legible and informative; see Figure 3). Dots show data for individual subjects, bars show the average response across subjects, and errors bars show standard errors of the mean across subjects. Thick, horizontal black lines are averaged across experiments, and gray rectangles are the corresponding 95% confidence intervals. Note that conditions from the same experiment share the same color; specifically, the six experiments that each contained both sentences and word lists conditions are presented at the end (right) of the Sentences bar group and the beginning (left) of the Word lists bar group, for east of comparison.

Critically, we also found a strong effect of task, such that responses in the MD fROIs were stronger in the experiments with an explicit task than in the passive reading/listening paradigms (b = 0.56, SE = 0.14, z = 4, p < 10^-4^); **Figure 5**; see **Figure SI-2** for responses of individual fROIs). In fact, some passive reading/listening experiments elicited a response at or below the fixation baseline in the MD network (**Figure SI-3)**. In contrast, in the language fROIs, the task did not affect the responses (b = −.18, SE = 0.14, z = −1.27, p = 0.203), with robust responses elicited by both experiments with an explicit task (b = 0.71, SE = 0.15, t(18.85) = 4.6, p < 10^-3^) and passive reading/listening paradigms (b = 0.89, SE = 0.16, t(22.38) = 5.5, p < 10^-4^). If anything, the latter elicited numerically stronger responses.

**Figure 5.**
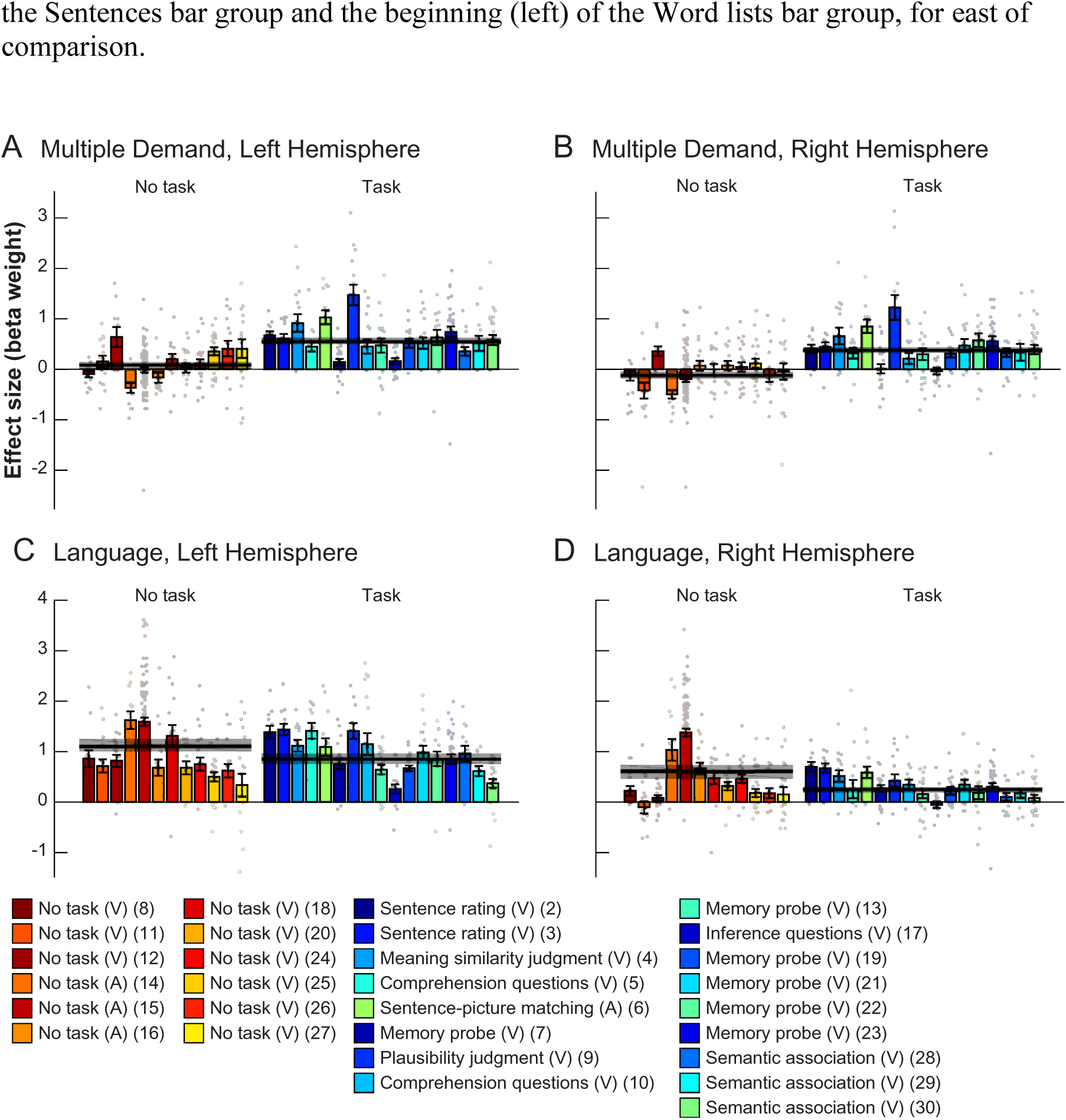
Responses of each network and hemisphere to passive and task-based **paradigms across experiments.** Same conventions as in Figure 4, but bars are now grouped by whether the experimental paradigm was passive comprehension or an active task. For experiments that contained both sentences and word lists, responses are averaged across these two conditions.

## Discussion

Across 30 fMRI language comprehension experiments (481 participants, 678 sessions), we examined how the brain regions of the domain-general Multiple Demand (MD) network (Duncan, 2010, 2013)—which have been linked to executive demands—respond to language processing. Consistent with prior work, we found above-baseline MD responses during many linguistic tasks. Moreover, these responses were stronger in the left hemisphere, mirroring the lateralization observed in the fronto-temporal language-selective network (Mahowald & Fedorenko, 2016). However, in sharp contrast to the language-selective network, which responds more strongly when participants process structured and meaningful stimuli (sentences) compared to individual words or lists of unconnected words, the MD network exhibited the opposite preference. This result already puts into question the key role of the MD network in any combinatorial linguistic operations—related to syntactic parsing or semantic composition—because one would expect brain regions that support such operations to respond more strongly to stimuli where those operations are engaged (i.e., sentences). But even more importantly, MD responses strongly depended on the presence of an explicit task: passive reading/listening tasks—which elicit strong responses in the language areas—elicited a much weaker response in the MD network (at the level of the fixation baseline, on average; **Figure 5A-B**) compared to experiments with a task.

Why might we, a priori, think that the domain-general MD network is important for language comprehension? There is a long tradition in the psycholinguistic literature to describe both lexical access and syntactic/semantic parsing using domain-general cognitive constructs. These include storing information in and retrieving it from working memory, updating focal attention, inhibiting irrelevant information, selecting an option among alternatives, and predictive processing (e.g., Johnson-Laird & Nicholas, 1983; Abney & Johnson, 1991; King & Just, 1991; Resnick, 1992; Gernsbacher, 1993; Waters and Caplan, 1996; Gibson, 1998; McElree, 2000, 2001; Gordon et al., 2002; Fedorenko et al., 2006, 2007; Lewis et al., 2006; Novick et al., 2009; Rodd et al., 2010; Schuler et al., 2010; Vergauwe et al., 2010; Smith & Levy, 2013; van Schijndel et al., 2013; Rasmussen & Schuler, 2018). These kinds of mental operations may be implemented in domain-general circuits of the MD network, which has historically been linked to diverse executive demands (Miller & Cohen, 2001; Duncan & Owen, 2000; Duncan, 2010). Indeed, prior neuroimaging studies have attributed core linguistic computations, like the ones above, to (parts of) the MD network (e.g., Thompson-Schill et al., 2002; Novick et al., 2005; Hirshorn & Thompson-Schill, 2006; Moser et al., 2007; Grindrod et al., 2008; January et al., 2009; Strijkers et al., 2019). However, alternatively, computations like inhibiting irrelevant information or predictive processing—albeit similar across domains—may be implemented in domain-specific cortices that store the relevant knowledge representations (Hasson et al., 2015).

Our results support the latter possibility and argue against the role of the MD network in core aspects of language comprehension. If a brain region supports a computation that is part and parcel of language understanding, this computation should be performed regardless of whether we are processing language passively or whether language processing is accompanied by a secondary task, like a memory-probe or comprehension-question task. This is exactly the pattern we observe in the language-selective network, which exhibits a task-independent response profile. However, the MD network’s response during passive comprehension tasks is, on average, at the level of the fixation baseline. These findings suggest that the MD network’s engagement likely reflects artificial task demands rather than language comprehension per se, and that all the core linguistic computations (cf. section 1 below) take place outside of MD areas, presumably in the language-selective areas (see Blank & Fedorenko, 2017, for converging evidence, which suggests that language, but not MD, regions “track” naturalistic linguistic input closely, and that the MD network’s computations are therefore unlikely to be related to the input features; also Shain, Blank, et al., 2020; Wehbe et al., submitted). Below, we raise four issues important to consider in light of the main conclusion we’re drawing here—that the MD network does not support core aspects of sentence comprehension.

1. ‘Core’ linguistic computations not tapped by our materials? We here construe core linguistic computations as computations that have to be engaged in order to extract a meaningful representation from the linguistic signal. By this definition, these operations should be engaged regardless of whether we are processing linguistic input passively, or whether we have to perform some additional task on the input. Core linguistic computations include computations related to lexical access and combinatorial processing (syntactic parsing and semantic composition), both of which strongly engage the fronto-temporal language network (e.g., Fedorenko et al., 2010, 2012, 2020; Bautista & Wilson, 2016). Might the materials used in the current study— across the 30 experiments—not tap some core comprehension-related computations? Aside from the cases discussed in section 2 below, the current set of materials is biased toward written language (26 of the 30 experiments use written materials) and does not include linguistic exchanges (e.g., dialogs or multi-person conversations). We therefore leave open the possibility that some linguistic operations engaged during auditory comprehension (e.g., prosody-related computations; e.g., Kristensen et al., 2013) or during the processing of socially interactive materials may drive the MD network.
2. Noisy language comprehension? The stimuli in the current study were clearly perceptible and well-formed. This differs from naturalistic comprehension scenarios, which are characterized by both low-level perceptual and higher-level linguistic noise (speakers make false starts/errors, etc.). Long prominent in speech perception research (Mattys et al., 2012), noise has recently permeated models of sentence interpretation (e.g., Ferreira et al., 2002; Levy et al., 2008; Gibson et al., 2013; Traxler, 2014). Prior fMRI studies of acoustically (e.g., Adank, 2012; Hervais-Adelman et al., 2012; Wild et al., 2012; Scott & McGettigan, 2013; Vaden et al., 2013; Peelle, 2018) and linguistically (e.g., containing syntactic errors; Kuperberg et al., 2003; Nieuwland et al., 2012; Mollica et al., 2020) noisy signals have reported activation in regions consistent with the topography of the MD network. So, the MD network may be important for coping with signal corruption. This may also be the underlying cause of MD regions’ responses during non-native (L2) language processing (e.g., Pliatsikas & Luk, 2016) because the representations of linguistic input are plausibly noisier in L2 speakers (Futrell & Gibson, 2017). That said, detection of errors in the input necessarily relies on the knowledge of the statistics of the relevant domain (language, in this case) and correction of noisy input relies on the knowledge of how noise operates (e.g., what kind of errors speakers are likely to make). Both of these kinds of knowledge are likely to be stored within the language-selective network, not the MD network (see also e.g., Shain, Blank, et al., 2020). Furthermore, in non-linguistic domains, the MD network shows increased activity during *any* more cognitively demanding condition, not only conditions with noisy input (e.g., Duncan & Owen, 2000; Crittenden & Duncan, 2012; Fedorenko et al., 2013; Hugdahl et al., 2015). As a result, the mechanisms that support error detection and correction during language comprehension, and the MD network’s contribution to processing noisy input remain to be characterized.
3. Other populations? The current study focused on neurotypical young adults. However, our brains are notoriously plastic, and tissue not previously engaged in some function can assume that function in addition to its original function(s) or via repurposing (e.g., Feydy et al., 2002; Cramer, 2008; Kleim, 2011). The MD network might be especially plastic in this way, given that it flexibly supports diverse behaviors and modulates its responses based on current task demands (e.g., Freedman et al., 2001; Cromer et al., 2010; Jackson et al., 2016; Kumano et al., 2016). Recent behavioral (e.g., Martin & Allen, 2008; Corbett et al., 2009; El Hachioui et al., 2014; Bonini & Radanovic, 2015; Villard & Kiran, 2016; Simic et al., 2017; Wall et al., 2017) and neuroimaging (e.g., Brownsett et al., 2014; Geranmayeh et al., 2016, 2017; Sims et al., 2016; Meier et al., 2016; Stockert et al., 2020) studies have begun to suggest a possible role for the MD network in recovery from aphasia (see Hartwigsen, 2018, for a review). Related evidence comes from increases in the MD network’s activity during language processing in aging (e.g., Wingfield & Grossman, 2006). However, whether the MD engagement is functionally important, or simply reflects greater processing demands, remains to be discovered.
4. Language production? The current study focused on comprehension. Might the MD network support core operations in language production? Executive processes have been implicated in both lexical access and syntactic planning based on behavioral (e.g., Alm and Nilsson, 2001; Roelofs and Piai, 2011; Strijkers et al., 2011, cf. Ivanova & Ferreira, 2017), neuroimaging (e.g., Indefrey and Levelt, 2004; Shuster and Lemieux, 2005; Alario et al., 2006; Troiani et al., 2008; Eickhoff et al., 2009; Wilson et al., 2009; Adank, 2012; Geranmayeh et al., 2012; Grande et al., 2012; Heim et al., 2012), and patient (e.g., Ziegler et al., 1997; Nestor et al., 2003; Coelho et al., 2012; Endo et al., 2013) evidence. Although language production presumably relies on the same knowledge representations as comprehension, the computational demands differ. For example, syntactic operations are obligatory for producing correct linguistic output, but may be foregone during comprehension (Bock, 1995). In addition, production is more demanding, and follows a developmental time-course that resembles that of executive functions (e.g., Hartshorne & Germine, 2015). As a result, the MD network may support some aspects of language production, although—as with comprehension—it will be important to dissociate core linguistic processes from extraneous task demands (Blanco-Elorietta & Pylkkanen, 2017).

To conclude, we have ruled out a set of hypotheses about the contributions of the domain-general MD network to language comprehension. In particular, we showed that MD areas only respond in comprehension experiments in the presence of a secondary task. We have consequently argued that the MD network is unlikely to support core linguistic computations that relate to lexical access, syntactic parsing, or semantic composition. However, we leave open the possibilities that the MD network (i) supports linguistic computations not engaged by the current materials, including operations related to the processing of noisy linguistic input, (ii) helps compensate for language loss after brain damage or in healthy aging, or (iii) supports core linguistic computations during language production.

## Author contributions

EF designed research. All authors performed research and analyzed data. IB created the figures. ED and EF wrote the paper with input from other authors.

## Acknowledgements

We would like to acknowledge the Athinoula A. Martinos Imaging Center at the McGovern Institute for Brain Research at MIT, and its support team (especially Steve Shannon and Atsushi Takahashi). We thank former and current EvLab members (especially Zach Mineroff, Brianna Pritchett, Dima Ayyash, and Zuzanna Balewski) for help with data collection, Alfonso Nieto-Castañon for help with the comparison between two preprocessing and analysis pipelines, Cory Shain and Ted Gibson for comments on earlier versions of the manuscript, and Swathi Kiran for discussions of the role of the MD network in language recovery in aphasia. EF was supported by NIH awards R00-HD057522, R01-DC016607, and R01-DC016950, a grant from the Simons Foundation to the Simons Center for the Social Brain at MIT, and support from the Department of Brain and Cognitive Sciences and the McGovern Institute for Brain Research at MIT.

## Notes

Conflict of interest: The authors declare no competing financial interests.

https://osf.io/pdtk9/

